# SARS-CoV-2 infection induces mixed M1/M2 phenotype in circulating monocytes and alterations in both dendritic cell and monocyte subsets

**DOI:** 10.1101/2020.10.09.332858

**Authors:** Matic Sanja, Popovic Suzana, Djurdjevic Predrag, Todorovic Danijela, Djordjevic Natasa, Mijailovic Zeljko, Sazdanovic Predrag, Milovanovic Dragan, Ruzic Zecevic Dejana, Petrovic Marina, Sazdanovic Maja, Zornic Nenad, Vukicevic Vladimir, Petrovic Ivana, Matic Snezana, Karic Vukicevic Marina, Baskic Dejan

## Abstract

Clinical manifestations of SARS-CoV-2 infection range from mild to critically severe. The aim of the study was to highlight the immunological events associated with the severity of SARS-CoV-2 infection, with an emphasis on cells of innate immunity. Thirty COVID-19 patients with mild/moderate symptoms and 27 patients with severe/critically severe symptoms were recruited from the Clinical Center of Kragujevac during April 2020. Flow cytometric analysis was performed to reveal phenotypic and functional alterations of peripheral blood cells and to correlate them with the severity of the disease. In severe cases, the number of T and B lymphocytes, dendritic cells, NK cells, and HLA-DR-expressing cells was drastically decreased. In the monocyte population proportion between certain subsets was disturbed and cells coexpressing markers of M1 and M2 monocytes were found in intermediate and non-classical subsets. In mild cases decline in lymphocyte number was less pronounced and innate immunity was preserved as indicated by an increased number of myeloid and activated dendritic cells, NK cells that expressed activation marker at the same level as in control and by low expression of M2 marker in monocyte population. In patients with severe disease, both innate and adoptive immunity are devastated, while in patients with mild symptoms decline in lymphocyte number is lesser, and the innate immunity is preserved.

## Introduction

Severe acute respiratory syndrome coronavirus 2 (SARS-CoV-2) continues to infect millions of people worldwide, causing coronavirus disease (COVID-19). The severity of reported symptoms for COVID-19 ranges from mild to critically severe having significant potential for fatal outcome. Previous studies have revealed a certain pattern of changes in biochemical and hematological parameters, while researches on immunopathology underlying COVID-19 are in progress. Currently, there is no wide agreement of the scientific and medical community about diagnostic, treatment and prognostic importance of immunological parameters for routine practice [1–3].

In COVID-19 patients inflammatory factors such as C – reactive protein (CRP) and erythrocyte sedimentation rate (ESR) are generally elevated, and CRP level, in general, positively correlates with the severity of the infection. High procalcitonin (PCT) level is a highly specific marker of the presence of bacterial infection and elevated levels of aspartate aminotransferase (AST), alanine aminotransferase (ALT), lactate dehydrogenase (LDH), creatine kinase (CK), D-dimmer and prothrombin time were proposed to be markers of severe disease [4]. The majority of studies conducted so far found that interleukin-6 (IL-6) serum concentrations positively correlate with exacerbation of disease after 7-14 days of onset of symptoms [5]. Other studies indicate that an increased number of neutrophils in combination with lymphopenia and consequent increase in neutrophil-to-lymphocyte ratio was the prognostic factor for severe cases [6]. The results of the first research on lymphocyte populations' change in severe COVID-19 cases indicated a decreased number of T lymphocytes, an increased number of naive T helper cells, and a decrease in memory T helper cells (1). Also, the number of CD8+T cells, B cells, and natural killer (NK) cells were substantially reduced in COVID-19 patients, particularly in severe cases [5–7]. In COVID-19 patients with severe pulmonary inflammation expression of NKG2 marker on NK cells and cytotoxic T lymphocytes were markedly increased and tend to correlate with functional impairment, indicating disease progression [8]. Although neutrophilia and impairment of lymphocyte number and function in COVID-19 patients have been well described, fewer data are available on dendritic cells and monocytes [9]. Two groups of authors separately pointed to alterations in the activation status and morphology of monocytes in severe cases. They identified forward scatter high (FCS-high) [10] and side scatter high (SSC-high) [11] populations of monocytes that secrete IL-6, IL-10 and TNF-α. One paper described the depletion of plasmacytoid dendritic cells in patients with severe disease [12].

Detailed analysis of immune parameters in COVID-10 patients and a better understanding of features of immune response underlying distinctive courses of the disease will improve diagnostics, the prognosis of disease outcome, as well as treatment strategies. Here, we present novel observations about changes in morphology and activation status of the cells of the innate immunity in COVID-19 patients, which seem to correlate with the severity of the infection.

## Patients and methods

### Patients/ Study design and participants

Fifty-seven cases of COVID-19 patients who were hospitalized in the Clinical Center of Kragujevac were recruited in this study during April 2020. Inclusion criteria were: adults of male or female gender (≥18 years old), SARS-CoV-2 infection confirmed by real-time polymerase chain reaction (RT-PCR) and hospitalization. COVID-19 patients were diagnosed according to the World Health Organization's (WHO) interim guidance [13]. Clinical condition severity was classified in four categories: a) mild; mild clinical symptoms of upper respiratory tract viral infection; b) moderate: present signs of pneumonia without need for supplemental oxygen; c) severe: fever or suspected respiratory infection with compromised respiratory function; and d) critically severe: worsening of respiratory symptoms with the necessity for mechanical ventilation. Our cohort of 57 COVID-19 patients consisted of 30 cases of mild/moderate disease (MD) and 27 cases of severe/critical disease (SD). Five healthy subjects with a negative RT-PCR test for coronavirus were included in the study as a control group. Ethics Committee of Clinical Center Kragujevac approved this study (Nr 01/20-405) and prior initiation written informed consent was obtained from every subject or the patients’ legal representative if he or she was unable to communicate e.g. sedated on mechanical ventilation, according to the Declaration of Helsinki of the World Medical Association.

### Data collection

The patients’ data were obtained from hospital medical records (electronic and paper version) for each study subject according to the modified case record rapid recommendation concerning the patients with Covid-19 infection of WHO (Global COVID-19 Clinical Platform: novel coronavirus (Covid-19) - rapid version. Geneva: World Health Organization, 2020. (https://apps.who.int/iris/rest/bitstreams/1274888/retrieve). The data for the following variables were collected: age, sex, medical history, symptoms, and signs of Covid-19, severity assessment, radiological imaging, and laboratory findings. Blood samples for laboratory tests and flow cytometry analysis were collected at admission, i.e. before any treatment. The following hematological parameters were examined: white blood cell count (WBC) with WBC differential count (neutrophils, eosinophils, basophils, lymphocytes, and monocytes), red cell blood count (RBC), hemoglobin concentration and platelet count (PLT). The measured biochemical parameters included creatinine (CRE), blood urea nitrogen (3), glycemia (GLY), albumin (ALB), aspartate aminotransferase (AST), alanine aminotransferase (ALT), lactate dehydrogenase (LDH), creatine kinase (CK), D-dimmer, C-reactive protein (CRP) and procalcitonin (PCT). Analysis of hematological parameters and blood biochemical assays were performed with commercial reagents and according to good laboratory practices at the hematology and clinical biochemistry departments of the hospital.

### Flow cytometry

Whole blood samples with anticoagulant from 5 healthy subjects, 10 patients with mild and 10 patients with severe disease were stained with anti-HLA-DR PE and ECD, anti-CD3 ECD, anti-CD4 PE, anti-CD8 FITC, anti-CD19 PC7, anti-CD14 FITC, anti-CD16 PC5, anti-CD15 PE, anti-CD57 FITC, anti-CD56 PE, anti-CD11c PCP, anti-CD123 PE, anti-CD83 PE, anti-CD38 PE, anti-CD23 ECD and isotype controls (all from Beckman Coulter) for 20 minutes in the dark at +4°C. Samples were analyzed on the flow cytometer Cytomics FC500 (Beckman Coulter). Data were processed by FlowJo V.10.

### Statistical analysis

Descriptive analysis of collected data and hypothesis testing of observed differences in measured variables were used. Shapiro-Wilk test was employed for the evaluation of normality data distribution. Independent groups t-test and analysis of variance (ANOVA) were used for comparison between groups. Two-sided *p*-values of less than 0.05 were considered statistically significant. Commercial statistical program SPSS (version 19.0, SPSS Inc., Chicago, IL) was used for data analysis.

## Results

### Baseline Characteristics of COVID-19 Patients

The median age of patients was 58 years and 35 patients were male (Table 1). Fever (75.4%), cough (70.2%), and fatigue (33.3%) were the most common symptoms upon admission. According to physical examination, 57.9% of patients had diminished breath sound and 42.1% had crackles. Radiological findings (standard chest X-ray) showed individual pneumonic foci in 50.9% of patients and interstitial changes in 36.8% ones. Hematological parameters and biochemical analysis of the whole patient cohort showed that increased granulocytes, glycemia, ALT, LDH, CK, D-dimmer, and CRP were the most typical findings, while oxygen saturation and blood pH were under normal values in the majority of patients (S1 Table). The values of laboratory parameters were significantly different among MD and SD cases (Table 2). WBC, granulocyte percent, LDH, CK, CRP, PCT, pCO_2_ were higher in SD patients, while the lymphocyte and monocyte counts, albumin levels, oxygen saturation, and potassium concentrations were lower among SD patients compared to MD ones.

**Table 1.** Clinical characteristics of COVID-19 patients

| *Characteristic* | *Patients* |
| --- | --- |
| **Age, median (min-max)** | 58 (23-88) |
| **Gender** |  |
| **Male, n (%)** | 35 (61.4) |
| **Female, n (%)** | 22 (38.6) |
| **Severity of disease** |  |
| **Mild/moderate, n (%)** | 34 (59.6) |
| **Severe/critically ill, n (%)** | 23 (40.4) |
| **Symptoms** |  |
| **Fever, n (%)** | 43 (75.4) |
| **Cough, n (%)** | 40 (70.2) |
| **Shortness of breath, n (%)** | 13 (22.8) |
| **Chest pain, n (%)** | 7 (12.3) |
| **Myalgia, n (%)** | 3 (5.3) |
| **Headache, n (%)** | 6 (10.5) |
| **Weakness, n (%)** | 19 (33.3) |
| **Sore throat, n (%)** | 7 (12.3) |
| **Loss of smell, n (%)** | 2 (3.5) |
| **Gastrointestinal symptoms, n (%)** | 9 (15.8) |
| **Physical examination** |  |
| **Normal breath sound, n (%)** | 9 (15.8) |
| **Diminished breath sound, n (%)** | 33 (57.9) |
| **High – pitched breath sounds, n (%)** | 3 (5.3) |
| **Crackles, n (%)** | 24 (42.1) |
| **Wheezing, n (%)** | 4 (7.0) |
| **Chest X-ray** |  |
| **Without active lesions, n (%)** | 9 (15.8) |
| **Prominent bronchovascular** | 3 (5.3) |
| **markings, n(%)** |  |
| **Interstitial changes, n (%)** | 21 (36.8) |
| **Individual pneumonic foci, n (%)** | 29 (50.9) |
| **Diffuse pneumonic foci, n (%)** | 16 (28.1) |
| **Ground-glass opacities, n (%)** | 1(1.8) |
| **Homogenous opacities, n (%)** | 2(3.5) |

**Table 2.** Haematological and serum biochemistry parameters in COVID-19 patients with mild and severe disease

| *Parameters* | *Mild* | *Severe* | *p* |
| --- | --- | --- | --- |
| **WBC (x10<sup>9</sup>/L)** | 5.4 ± 1.7 | 9.7 ± 4.9 | <0.0001 |
| **Granulocytes (%)** | 67.2 ± 9.4 | 82.4 ± 8.6 | <0.0001 |
| **Lymphocytes (%)** | 21.1 ± 6.7 | 9.3 ± 6.6 | <0.0001 |
| **Monocytes (%)** | 10.5 ± 3.5 | 6.7 ± 3.3 | <0.0001 |
| **AST(IU/L)** | 32.3±15.0 | 54.7±31.2 | 0.017 |
| **Albumins (g/L)** | 36.6 ± 4.3 | 29.6 ± 6.4 | <0.0001 |
| **LDH (U/L)** | 450.8 ± 99.5 | 919.2 ± 282.4 | <0.0001 |
| **CK (U/L)** | 124.7 ± 132.0 | 210.8 ± 144.6 | 0.035 |
| **CRP (mg/L)** | 36.3 ± 41.5 | 119.3 ± 80.9 | <0.0001 |
| **PCT (ng/mL)** | 0.1 ± 0.1 | 0.3 ± 0.3 | 0.001 |
| **pCO<sub>2</sub> (kPa)** | 4.7 ± 0.6 | 6.0 ± 3.1 | 0.048 |
| **Saturation (%)** | 96.1 ± 2.4 | 89.1 ± 6.7 | <0.0001 |
| **K(mmol/L)** | 4.4 ± 0.5 | 3.9 ± 0.6 | 0.003 |

The numbers represent the means ± standard deviations

### Changes in the frequency of peripheral blood cells in COVID-19 patients

Both mild and severe cases had a higher percent of polymorphonuclear, but lower percentage ratio of mononuclear cells in comparison to healthy controls, and that difference was more profound in severe cases (Table 3.).

**Table 3.**
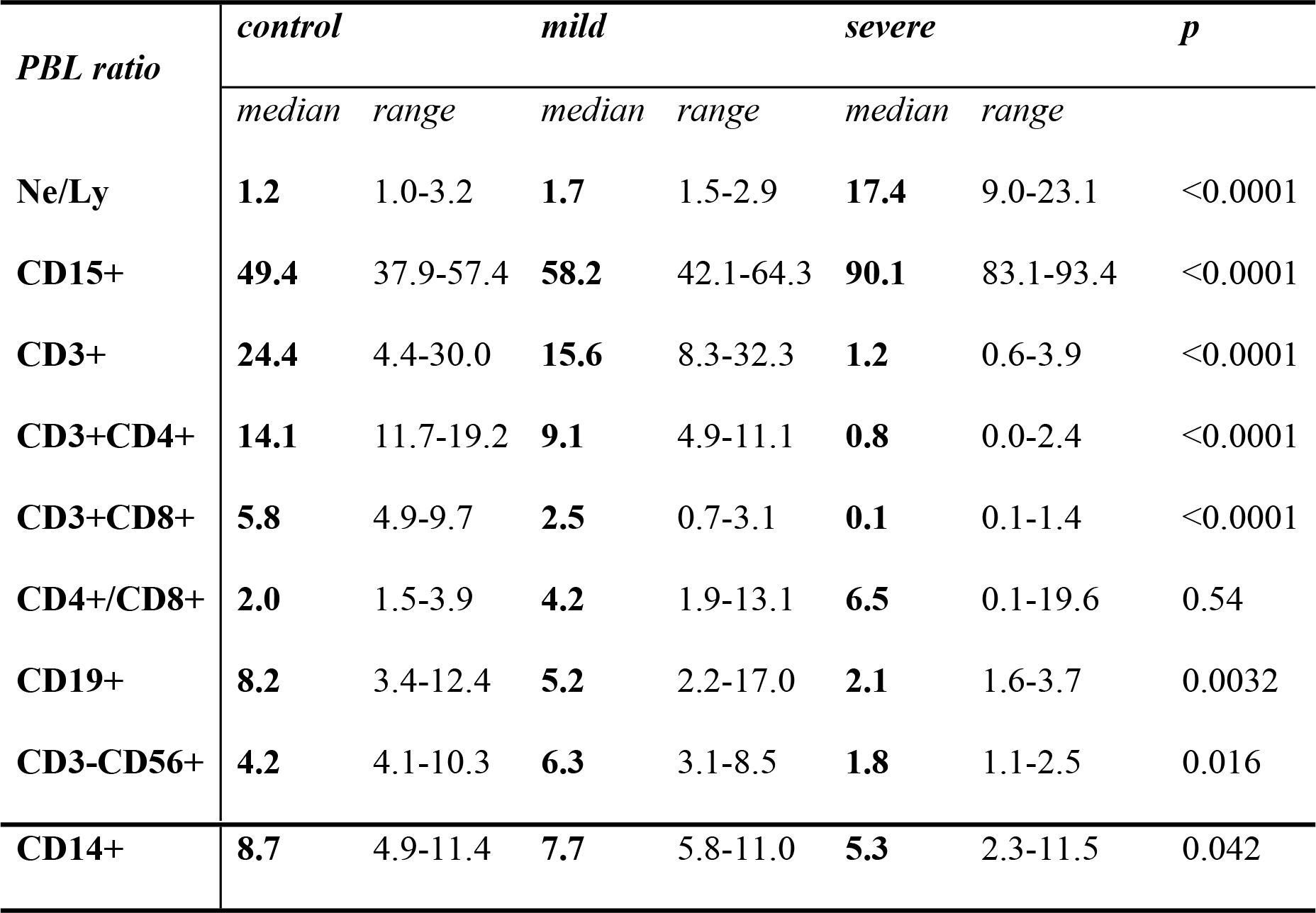
The percentage ratio of peripheral blood leukocytes (PBL) in healthy subjects, patients with mild disease, and patients with severe disease, as shown by flow cytometric analysis.

Although in non-severe patients all parameters were close to, or in the normal range, percent of CD15+ cells (neutrophils) was higher than in controls, and proportions of B lymphocytes (CD19+), monocytes (CD14+) and both helper (CD3+CD4+) and cytotoxic (CD3+CD8+) T lymphocytes were lower. In patients with severe disease very high neutrophil-to-lymphocyte ratio (17.4) reflected an increase in neutrophil count (90.1%), diminution in B lymphocyte (2.1%) and NK cells count (1.8%), and marked decline of T lymphocyte percentage (1.2%). Very low percent, far below the lower limit of the normal range, was found for both helper (0.8%) and cytotoxic (0.1%) T cells. CD4/CD8 ratio was three times higher in severe patients than in controls (S1 Fig), but without statistical significance. The percentage of monocytes/macrophages was in the normal range in both mild and severe cases, although lower than in control.

The NK cell subpopulation (CD3-CD56+) was assessed for the expression of CD57, a marker of NK cell maturation and activation. The change in NK cell number was statistically significant among the groups (*p*=0.016). In relation to control, mild cases had a higher number of NK cells (6.3% vs. 4.2% in controls), but about the same percent of CD57+ cells (1.1% in mild cases; 1.2% in controls), while in severe cases both total number of NK cells and percent of activated cells were lower (1.8% and 0.5%, respectively) (Fig 1), even though statistical significance was not reached.

**Fig 1.**
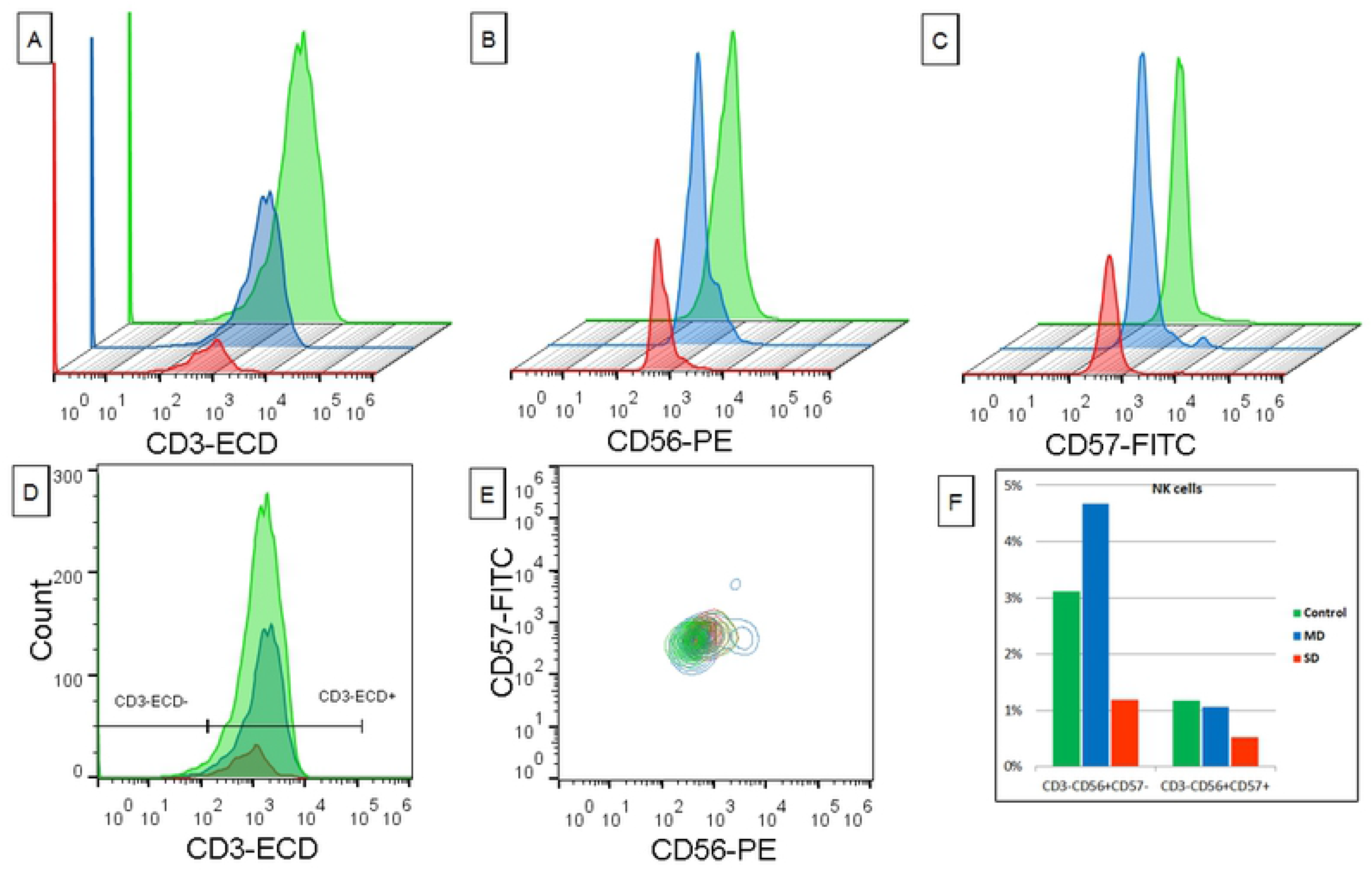
Flow cytometry analysis of NK cells in the whole blood of COVID-19 patients. (A-C) Smoothed histograms: CD3, CD56 and CD57 expression in patients with severe disease (red), mild disease (blue) and healthy control (green). (D) Overlaid histograms: Gating strategy for NK cells. (E) Overlaid contour plot: Identification of CD3^-^CD56^+^CD57^+^ cells in patients with severe disease (red), mild disease (blue) and healthy control (green). (F) Bar chart: The percentage of CD3^-^CD56^+^CD57^-^ and CD3^-^CD56^+^CD57^+^ cells in healthy controls and patients with mild (MD) and severe disease (SD).

Further analysis demonstrated that the percent of cells expressing HLA-DR was almost two times lower in mild cases than in controls (8.1% vs. 14.9%), and 6.5 times lower in severe cases (2.3%) with statistical significance of *p*<0.0001. Namely, a decrease of HLA-DR expression, more pronounced in severe cases, was determined in both monocytes (5.5% - controls; 4.2% - mild cases; 1.5% - severe cases, *p*<0.0001) and B lymphocytes (2.0% - controls; 1.3% - mild cases; 0.7% - severe cases, *p*=0.014) (Fig 2).

**Fig 2.**
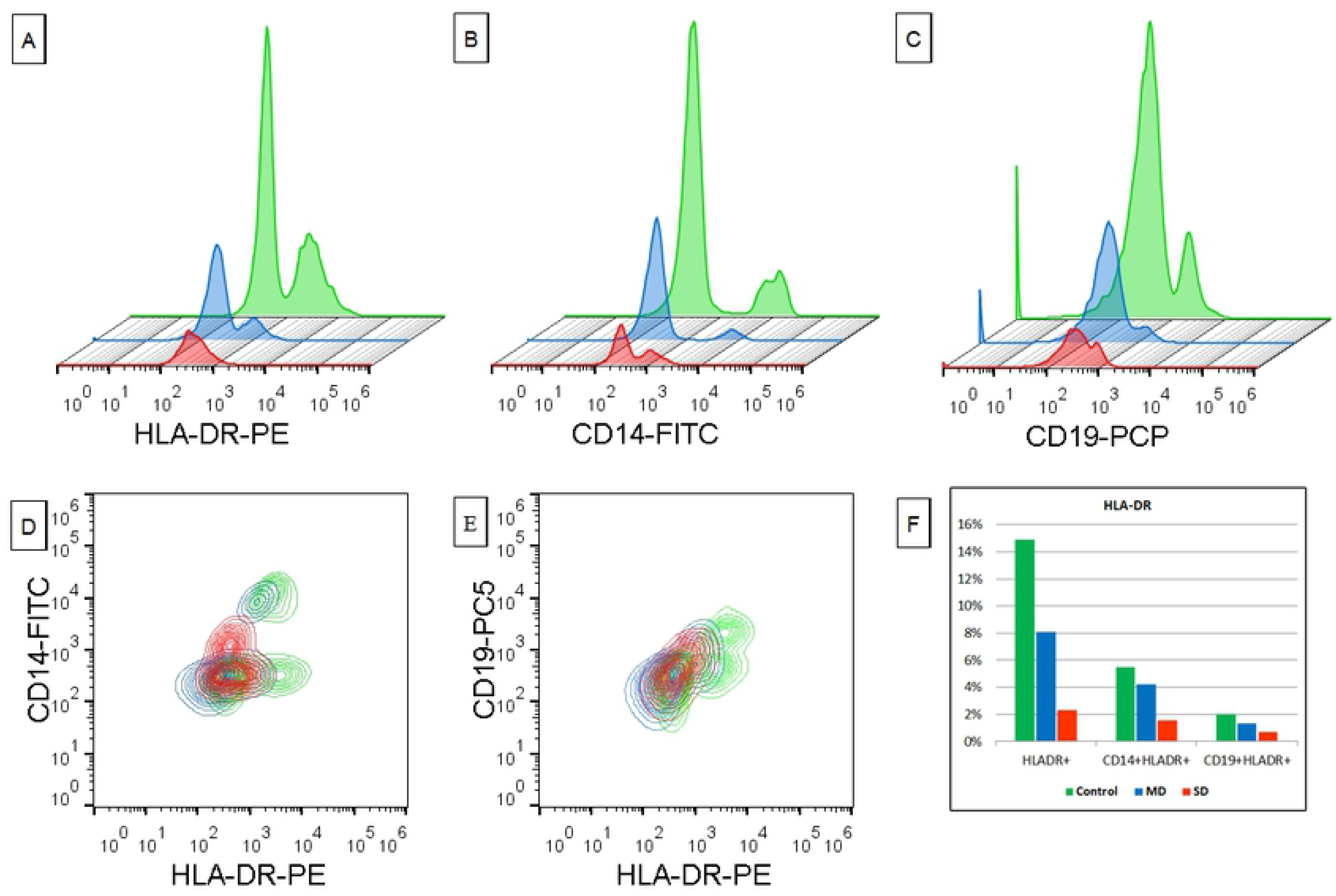
Flow cytometry analysis of the HLA-DR, CD14 and CD19 expression in whole blood of COVID-19 patients. (A-C) Smoothed histograms: HLA-DR-PE, CD14-FITC and CD19-PC5 expression showing patients with severe disease (red), mild disease (blue) and healthy control (green). (D, E) Dot plots: Identification of HLA-DR^+^CD14^+^ and HLA-DR^+^CD19^+^ population with color representing patient with severe disease (red), mild disease (blue) and control (green). (F) Bar chart: The percentage of HLA-DR^+^, HLA-DR^+^CD14^+^ and HLA-DR^+^CD19^+^ expression in healthy controls and patients with mild (MD) and severe disease (SD).

The percentage ratio of dendritic cells (Lyn-HLADR+) didn’t differ much between controls and mild cases, but in later, there was a lower number of plasmacytoid (CD123+) DCs (0.6% vs. 1.4% in control, *p*=0.0017), a higher number of myeloid (CD11c+) DCs (1.0% vs. 0.2% in control, *p*=0.0047), and DCs expressing CD83, an activation marker for antigen-presenting cells (0.7% vs. 0.5% in control) (Fig 3). Contrarily, in severe cases, CD11c+ DCs were almost undetectable and the percent of activated and plasmacytoid DCs was lower than in mild cases and controls (0.1%).

**Fig 3.**
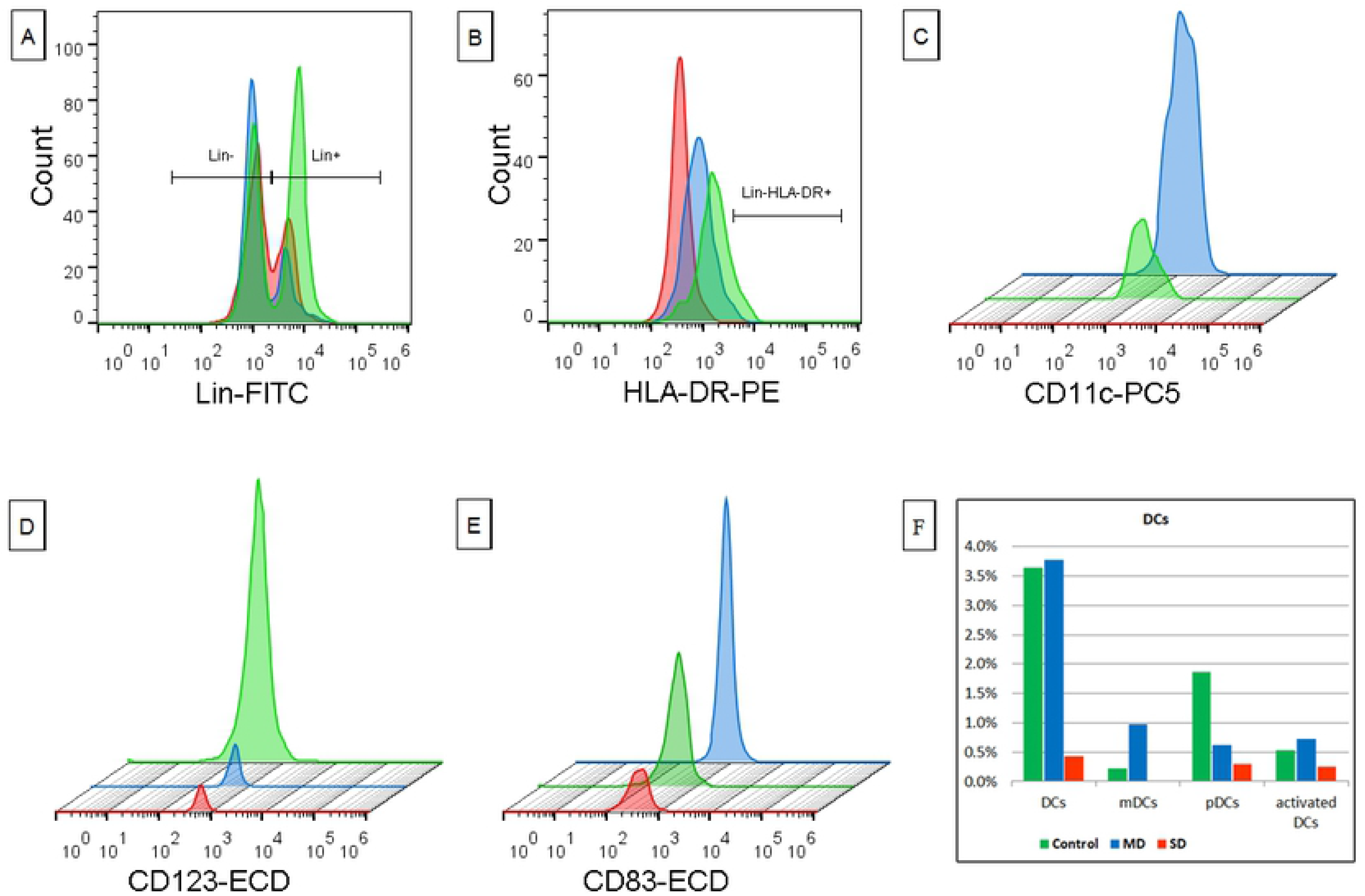
Flow cytometry analysis of dendritic cells in the whole blood of COVID-19 patients. (A-B) Overlaid histograms: Gating strategy for DC. (C-E) Smoothed histograms: CD11c, CD123 and CD83 expression in dendritic cells (Lin^-^HLA-DR^+^) of patients with severe disease (red), mild disease (blue) and healthy control (green). (F) Bar chart: The percentage of myeloid (mDCs), plasmacytoid (pDCs) and activated dendritic cells in healthy controls and patients with mild (MD) and severe disease (SD).

Further, by tracking relative expression levels of CD14 and CD16 surface molecules, we examined the proportions of monocyte/macrophage subsets, classical, intermediate, and non-classical (S2 Fig.). In relation to control the percent of classical monocytes (CD14^high^CD16-) was higher in mild cases and even higher in severe cases with statistical significance (86.0% in control, 89.2% in mild cases and 94.8% of total monocytes in severe cases, *p*=0.0033), while the percent of intermediate (CD14^high^CD16+) and non-classical monocytes (CD14^low^CD16+) decreased (Fig 4) (intermediate: 5.5% in control, 3.6% in mild and 1.9% in severe cases, *p*=0.14; non-classical: 8.6% in control, 7.2% in mild and 3.3% in severe patients, *p*=0.0057).

**Fig 4.**
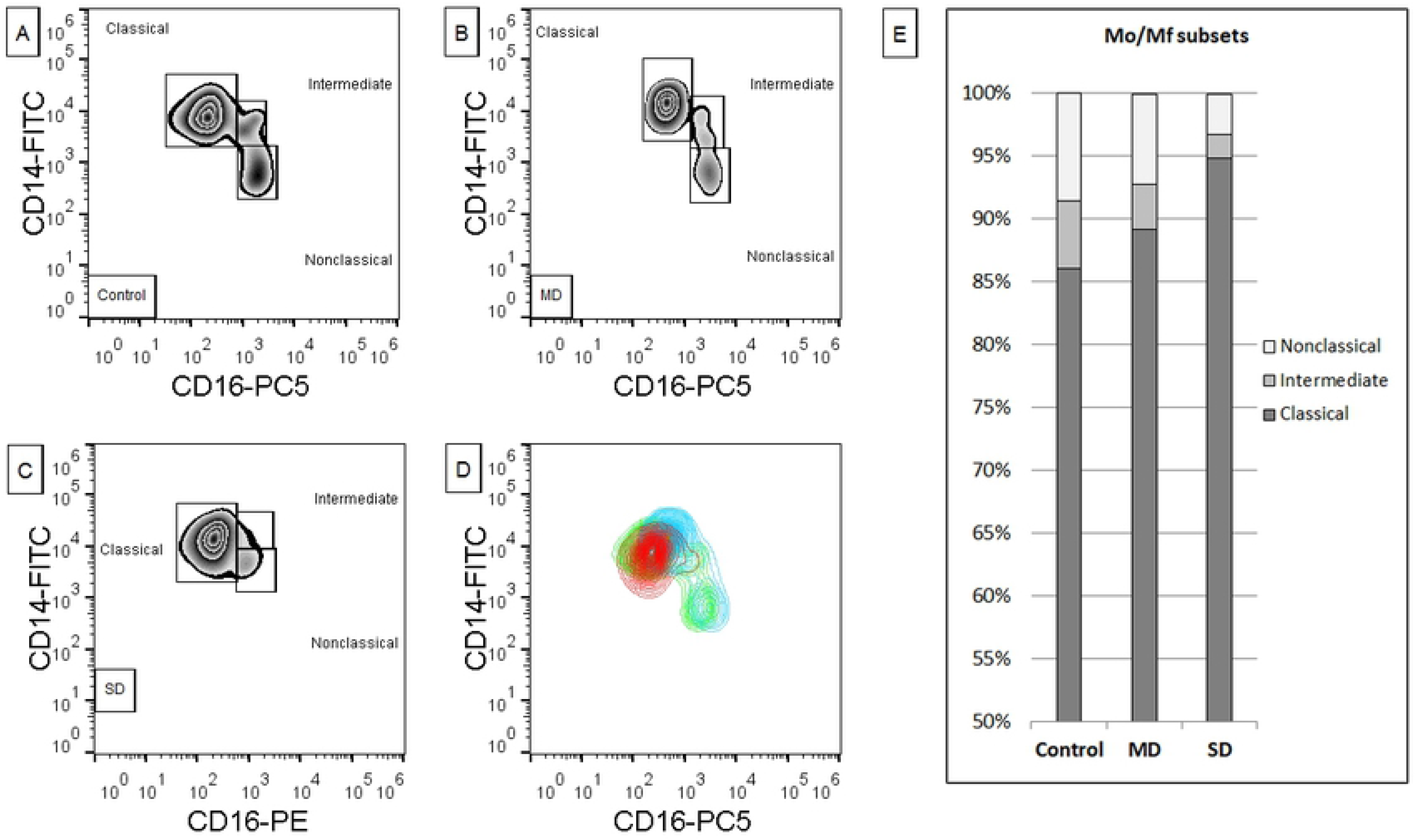
Flow cytometry analysis of monocyte subsets. (A-C) Zebra plots: Monocyte subsets in healthy control, patients with mild (MD) and severe disease (SD). (D) Overlaid contour plot: Monocyte subsets in patients with severe disease (red), mild disease (blue) and healthy control (green). (E) Bar chart: The percentage of the classical, intermediate and non-classical monocytes in healthy controls and patients with mild (MD) and severe disease (SD).

Next, the polarization of monocytes was revealed using CD38 as a marker of M1 monocytes, and CD23 typical of M2 monocytes. In the monocyte population of severe cases we found higher percent of CD38+ (98.3% vs. 94.1% in control and 93.3% in mild cases, *p*=0.039) and a markedly higher percent of CD23+ cells (10.1% vs. 1.5% in control and 1.6% in mild cases, *p*=0.0032). Of note, all monocytes positive for CD23, co-expressed CD38 as well (Fig 5).

**Fig 5.**
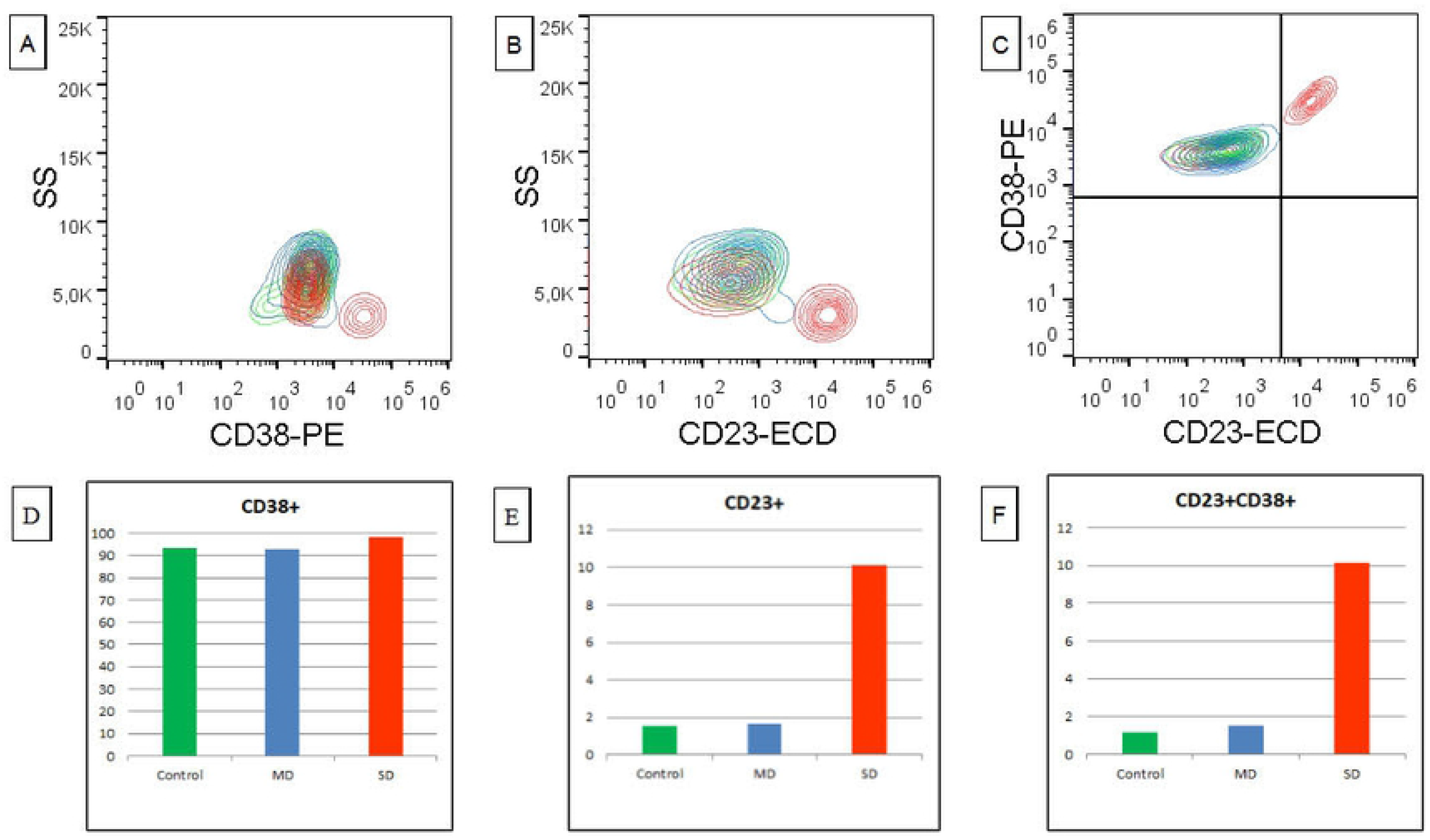
Flow cytometry analysis of the CD38 and CD23 expression in monocytes of COVID-19 patients. (A-C) Overlaid contour plot: Identification of CD38^+^, CD23^+^ and CD38^+^CD23^+^ monocytes in patient with severe disease (red), mild disease (blue) and control (green). (D-F) Bar chart: The percentage of the CD38^+^, CD23^+^ and CD38^+^CD23^+^ monocytes in healthy controls and patients with mild (MD) and severe disease (SD).

Cells co-expressing CD23 and CD38 were absent in classical monocyte subset of both controls and patients, but they were present in intermediate and non-classical subsets in patients, especially in severe cases (intermediate: 5.9% - control, 21.5% - mild, 27.7% - severe cases, *p*=0.17; non-classical: 3.6% - control, 17.1% - mild, 48.5% - severe cases, *p*=0.0021) (Fig 6).

**Fig 6.**
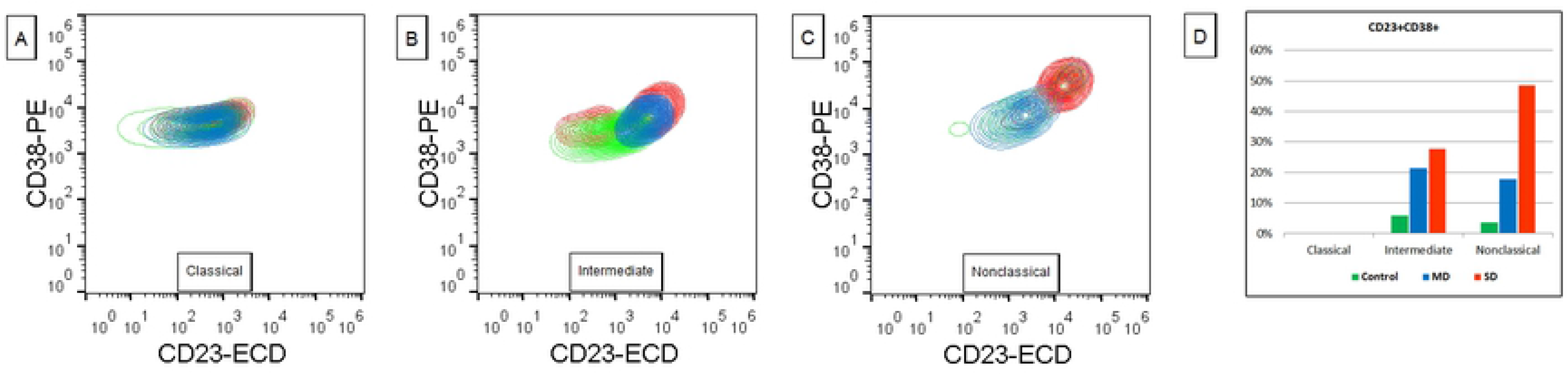
Flow cytometry analysis of the CD38 and CD23 expression in monocyte subsets of COVID-19 patients. (A-C) Overlaid contour plot: Identification of CD38^+^CD23^+^ in classical, intermediate and non-classical monocytes of the patient with severe disease (red), mild disease (blue) and control (green). (D) Bar chart: The percentage of the CD38^+^CD23^+^ classical, intermediate and non-classical monocytes in healthy controls and patients with mild (MD) and severe disease (SD).

## Discussion

COVID-19 is the third emerging coronavirus infectious disease in the 21st century. The virus was introduced from an animal reservoir and met an immunologically naive human population. A number of studies have described changes within innate and adaptive immune response in SARS-Cov-2-infected patients, but there are still many unknowns. In our study, we recorded baseline characteristics of COVID-19 patients and analyzed changes in peripheral blood cell populations in relation to control healthy subjects.

SARS-Cov-2 infection provokes sustained cytokine and chemokine secretion, leading to severe lung injury, multiorgan failure, immune dysfunction, and mortality. Our results are in line with previous findings showing that these alterations came under the signs of a common respiratory infection such as fever, cough, and fatigue [14] and that most patients had granulocytosis, elevated infection-related and organ-injuries-related biomarkers, including LDH, CK, ALT, which were higher among severe cases. Also, we found elevated levels of D-dimmer and CRP, parameters that have been reported to be associated with the severity of disease [4]. The cytokine storm is thought to be responsible for multiorgan damage and elevated organ-injuries-related biomarkers. However, correlation analysis between lymphocyte subsets and biochemical markers showed that most biochemical markers indicating organ damage are negatively correlated with lymphocyte counts in SARS-Cov-2 patients, which is not the case in patients with pneumonia of other etiologies. This finding highlights that the potential cause of multi-organ injury is the virus itself [15].

The common findings of previous research were high percent of neutrophils, lymphopenia, and high neutrophil-to-lymphocyte ratio in COVID-19 patients in comparison to healthy subjects, whereas this difference was more radical in severe disease cases, which is in consent with our results [6, 16-19]. Lymphopenia is one of the most salient markers of COVID-19 and it seems to arise as the result of the reduction of all lymphocyte populations, including CD4+ and CD8+ T cells, B cells, and NK cells. In line with our results, Zhou et al. [18] have described that the decline in CD4+ T lymphocyte count is significant in both severe and mild patients, while the decrease of CD+ 8 cells was more profound in severe patients. Still, there is a report that the reduction in CD4+ T cells is much greater in severe cases [6]. As previously described, we found a greater decrease in CD8+ than in CD4+ subpopulation, and the decrease in B cell percentage, that was more expressed in severe cases. It’s noteworthy that the decline in the frequency of all lymphocyte populations is more profound in COVID-19 patients in comparison with non-SARS-Cov-2-pneumonia patients [15]. One of the possible causes of lymphopenia is the sequestration of lymphocytes in the lung tissue, at the site of infection. The autopsy showed that infiltrating cells were mostly monocytes and macrophages, with multinucleated giant cells, and a few lymphocytes, being mainly CD4+ T cells [20]. Further examination also revealed that the number of trilineage in the bone marrow and lymphocytes in the spleen and lymph nodes are all significantly reduced. These facts indicated that lymphopenia cannot be attributed only to the tissue redistribution of lymphocytes and brought to the foreground the possible direct effect of the virus on immune cells. MERS-CoV is known to directly infect human T lymphocytes and activate the extrinsic or intrinsic apoptotic pathway, but does not replicate. Although ACE2 is not expressed on lymphocytes, as the main site of SARS-Cov-2 binding, recent research has reported a novel invading route of SARS-Cov-2. Namely, it has been noted that SARS-Cov-2 can infect lymphocytes through spike protein (SP) interaction with lymphocyte’s CD147 protein, highly expressed on activated T and B cells, but also on dendritic cells, monocytes and macrophages [21–22]. Whether virus induces direct cytopathic effects is not yet elucidated.

NK cells, as a member of innate immunity, provide crucial early defense against viral infections. Contrarily to other reports [7, 8, 16], our results showed that in mild COVID-19 patients the percentage of total NK cells was higher in comparison to control, and the percent of activated cells was preserved, but in severe cases, both values were remarkably decreased. It has been described that type I IFN is required for NK cell activation [23]. Low percent of dendritic cells in severe patients, as described, may contribute to decreased secretion of type 1 IFN, resulting in a decline of NK cell activation. Further, the overproduction of IL-6 plays a role in the reduced activity of NK cells in mimicked viral infection *in vivo* [24]. Recent findings state that IL-6 level is high in severe COVID-19 patients, remaining low in mild cases [5], and negatively correlates with NK cell count and activity [7]. Moreover, some authors noted upregulated expression of the inhibitory receptor NKG2A on NK cells in the early stage of COVID-19 [8].

Abnormalities in cells that bridge innate and adaptive immunity, and regulate the latter may be responsible for the reduction of lymphocyte number and function. Key roles in immune response regulation and antigen presentation play dendritic cells (DCs). A number of ways in which viruses affect the adequate response of DCs have been described [25]. In our study, we found that the percentage of overall DCs, as well as the percent of myeloid and activated DCs, was higher in mild cases in relation to control, indicating their preserved antigen-presenting function. Also, Cao et al. [26] have demonstrated that in Influenza virus-induced differentiation of monocytes into mDCs, those cells, unlike classic mDCs, secrete chemoattractants for monocytes and type I IFN. Contrarily, in severe cases, the percent of total DCs and analyzed subpopulations was lower. Functional activation analysis of DCs in SARS-CoV infection were inconclusive showing both activation [27], and the absence of activation [28]. The consequence of low degree dendritic cell activation is the insufficiency of costimulatory molecules, necessary for survival during TCR engagement, which partly explains the reduction of T lymphocytes dying by apoptosis in the absence of adequate signaling. Reduced percentage of plasmacytoid dendritic cells that we found in both mild and severe patients, suggests that the adequate response to viral infection was profoundly disrupted, considering that plasmacytoid DCs represent the main source of type I IFN. A similar effect was observed in SARS-CoV infection, where DCs failed to trigger a strong type I IFN response, implicating that the virus circumvents the activation of the innate immune system [29]. SARS-CoV also promoted a moderate increase in the production of IL-6 in DCs [30].

The expression of HLA-DR molecules is restricted to the cells with a specialized role in antigen presentation. Therefore, the extent of HLA-DR expression on monocytes and B cells indicates their ability for antigen-presentation. In our study overall expression of HLA-DR molecule on peripheral blood mononuclear cells was extremely downregulated among both mild and severe COVID-19 patients. The decline in monocyte HLA-DR expression in SARS-CoV-2 infection has been described in recent studies [11, 19], pursuant to our results. It should be underlined that we also found reduced HLA-DR expression on B lymphocytes. Of note, in our cohort HLA-DR expression was even 6.5 times lower in severe patients. Suppression of HLA-DR expression on monocytes below the threshold value (<30% HLA-DR + monocytes) has been accepted as a definition of immunoparalysis that occurs in lethal conditions such as sepsis and represents a predisposition for superinfection with a variety of pathogens [31]. Giamarellos-Bourboulis et al. [19] proposed that one of the drivers of decreases in HLA-DR expression is IL-6 concentration, based on finding that IL-6 concentration is reciprocal to HLA-DR expression.

The percentage ratio of monocytes in COVID-19 patients was in the normal range, but flow cytometric analysis showed that they are different from those in healthy subjects. Although we didn’t find FSC-high and SSC-high monocyte population described by some authors [10, 11], we revealed that the proportion between certain subsets of monocytes was disturbed in patients in comparison to controls. In mild cases classical (CD14^high^CD16-) monocytes were presented in higher percent, whereas the ratio of intermediate (CD14^high^CD16+) and non-classical (CD14^low^CD16+) decreased. This difference was more pronounced in severe cases. In the peripheral blood of healthy humans, classical monocytes are the major population of monocytes (80-95%) [32]. The main function of these so-called “inflammatory” monocytes is phagocytosis and secretion of proinflammatory cytokines. Besides, they are the primary source of monocyte-derived DCs and tissue macrophages. Intermediate monocytes (2-8%), which also have inflammatory properties, are the main ROS producers and have the highest expression of MHC II class molecules (HLA-DR), acting as specialized antigen-presenting cells. Non-classical monocytes (2-11%) are “patrolling” monocytes that travel across blood vessels to scavenge dead cells and pathogens. In contact with infectious agents, they produce inflammatory cytokines and chemokines that recruit neutrophils, and subsequently clear resulting debris and promote healing and tissue repair. During inflammation, non-classical monocytes can extravasate to inflamed tissue and differentiate to inflammatory macrophages. The three monocyte subsets have distinct roles in response to different stimuli, i.e. during homeostasis, inflammation, and tissue repair. In infectious diseases, the ratio of monocyte subsets varies depending on the pathogen, but an augmentation of CD16+ monocytes was found in the majority of cases. Prominent increase of CD16+ monocytes was determined in patients with bacterial infections and bacterial sepsis [33]. An increase in the percent of non-classical monocytes was observed during HIV [34], Hepatitis C [35], and Dengue virus infections [36]. As well, an increase in CD16+ monocytes was reported in COVID-19 patients by Zhang and Zhou [10, 18], while Hussman described raise in the intermediate subset in patients with severe pulmonary complications [37]. In contrast to these reports, we found an augmentation of classical (CD16-) subset and concomitant diminution of CD16+ subsets in COVID-19 patients. This may be a result of the migration of non-classical monocytes into the lungs where they differentiate and become inflammatory macrophages.

To further define functional changes in monocyte subsets during infection, we observed the expression of surface molecules CD38 and CD23. Our results showed a high expression of both CD38 and CD23 on monocytes in patients with severe disease. Importantly, in COVID-19 patients monocytes co-expressing CD23 and CD38 were found in intermediate and nonclassical subsets. Mixed M1/M2 phenotype has been described in chronic infections, autoimmune diseases, cancer, and disorders associated with fibrosis (38). It is postulated that these monocytes have both proinflammatory and tissue repair functions. A decrease in the number of nonclassical monocytes in COVID-19 patients can be explained by their extravasation to lungs, where M1/M2 cells promote both inflammation and fibrosis as a tissue repair mechanism. Pulmonary fibrosis, a characteristic of acute respiratory distress syndrome (ARDS), is a complication of COVID-19 and can also be a cause of mortality in COVID-19 patients [39]. Considering the central role that monocytes play in the pathogenesis of cytokines storm in lung damage of COVID-19, here we present both conformation of some already available evidence and add some novelty about substantial phenotypical alterations of monocytes in COVID-19 patients.

Overall, pursuant to the previous findings and based on our results, we can speculate that SARS-CoV-2 virus causes dismantling of the immune response, but the resulting alterations differ in mild and in severe patients. In patients who didn't develop severe symptoms, a decline in lymphocyte number is lesser, and the innate immunity is preserved. Despite a decrease in the percent of HLA-DR-expressing B cells and monocytes, the number of total DCs, mDCs (possibly type I IFN producing cells), and activated DCs is higher than in control, pointing to their sustained functions. NK cells, that play a key role in host defense against viral infections, are also present in higher number, and activated cells are immanent as in healthy subjects. In monocyte population expression of M2 marker CD23 is low, as in healthy controls. On the other hand, in severe cases, both arms of immune defense, innate and adoptive, are affected. The number of T and B lymphocytes is dramatically decreased, as well as the number of NK cells and DCs (both total and activated). The number of cells expressing HLA-DR is drastically lesser in severe cases, depicting the inability of antigen-presenting cells to activate T lymphocyte response. The virus affects the monocyte population as well. The percentage ratio of intermediate and nonclassical monocytes is reduced, pointing to impaired functional maturation. An increase in the percent of cells coexpressing markers of M1 and M2 monocytes points to prolongation of inflammation and evolution of fibrosis as a repair mechanism, that damages lung parenchyma and potentially increases the risk of worse clinical outcomes. Altogether, our results depict the devastation of host defense in severe patients and altered, but a more efficient immune response in patients with mild/moderate symptoms.

## Acknowledgments

We thank the health workers of the Clinical Centre Kragujevac, doctors and especially technicians, who selflessly, in difficult moments of the struggle for the lives of patients, supported our study.

## Supporting information captions

**S1 Table. Haematological and serum biochemistry parameters in COVID-19 patients.** The numbers represent the means ± standard deviations

**S1 Fig. Flow cytometry analysis of T cells in whole blood of COVID-19 patients.** (A-C) Smoothed histograms: CD3, CD4 and CD8 expression in patients with severe disease (red), mild disease (blue) and healthy control (green). (D, E) Overlaid contour plot: Identification of CD3^+^CD4^+^ and CD3^+^CD8^+^ T cells in patients with severe disease (red), mild disease (blue) and healthy control (green).

**S2 Fig. Monocyte gating strategy in whole blood.** (A) FS *vs.* SS plot: Wide selection of monocytes depending on FS/SS properties. (B) Pseudocolor CD16 *vs.* CD14 plot: Gating to select monocytes depending on characteristic “inverted L” shape. (C) Pseudocolor CD16 *vs.* HLA-DR plot: Gating to select HLA-DR^+^ cells and to remove NK cells. (D) Pseudocolor CD14 *vs.* HLA-DR plot: Gating to exclude B cells (HLA-DR^high^/CD14^low^). (E) Pseudocolor CD16 *vs.* CD14 plot: Gating to select classical (CD14^high^CD16^-^), intermediates (CD14^high^CD16^+^) and non-classical (CD14^low^CD16^+^) monocytes. (F) Zebra CD16 *vs.* CD14 plot: Selected monocytes redisplayed on CD16 vs. CD14 zebra plot to visualize monocyte subsets.

**S3 Dataset**

## References

1. Lin L, Lu L, Cao W, Li T. Hypothesis for potential pathogenesis of SARS-CoV-2 infection-a review of immune changes in patients with viral pneumonia. Emerg Microbes Infect. 2020; 9(1): 727–732. doi:10.1080/22221751.2020.1746199

2. Felsenstein S, Herbert JA, McNamara PS, Hedrich CM. COVID-19: Immunology and treatment options. Clin Immunol. 2020; 215: 108448. doi:10.1016/j.clim.2020.108448

3. Chen R, Sang L, Jiang M, Yang Z, Jia N, Fu W, et al. Longitudinal hematologic and immunologic variations associated with the progression of COVID-19 patients in China. J Allergy Clin Immunol. 2020; S0091–6749(20)30638-2. doi:10.1016/j.jaci.2020.05.003

4. Ying-Hui J, Lin CA, Zhen-Shun C, Hong C, Tong D, Yi-Pin F, et al. A rapid advice guideline for the diagnosis and treatment of 2019 novel coronavirus (2019-nCoV) infected pneumonia (standard version). Mil Med Res. 2020; 7(1): 4. Published 2020 Feb 6. doi:10.1186/s40779-020-0233-6

5. Wan S, Yi Q, Fan S, Lv J, Zhang X, Guo L, et al. Characteristics of lymphocyte subsets and cytokines in peripheral blood of 123 hospitalized patients with 2019 novel coronavirus pneumonia (NCP). medRxiv 2020.02.10.20021832; doi: https://doi.org/10.1101/2020.02.10.20021832

6. Qin C, Zhou L, Hu Z. Dysregulation of immune response in patients with COVID-19 in Wuhan, China. Clin Infect Dis. 2020; ciaa248. doi:10.1093/cid/ciaa248

7. Wang F, Nie J, Wang H, Zhao Q, Xiong Y, Deng L, et al. Characteristics of Peripheral Lymphocyte Subset Alteration in COVID-19 Pneumonia. J Infect Dis. 2020; 221(11): 1762–1769. doi:10.1093/infdis/jiaa150

8. Zheng M, Gao Y, Wang G, Song G, Liu S, Sun D, et al. Functional exhaustion of antiviral lymphocytes in COVID-19 patients. Cell Mol Immunol. 2020; 17(5): 533–535. doi:10.1038/s41423-020-0402-2

9. Spiegel M, Schneider K, Weber F, Weidmann M, Hufert FT. Interaction of Severe Acute Respiratory Syndrome-Associated Coronavirus With Dendritic Cells. J Gen Virol. 2006; 87(Pt 7): 1953–60. doi: 10.1099/vir.0.81624-0

10. Zhang D, Guo R, Lei L, Liu H, Wang Y, Wang Y, et al. COVID-19 infection induces readily detectable morphological and inflammation-related phenotypic changes in peripheral blood monocytes, the severity of which correlate with patient outcome. medRxiv 2020.03.24.20042655; doi: https://doi.org/10.1101/2020.03.24.20042655

11. Lombardi A, Trombetta E, Cattaneo A, Castelli V, Palomba E, Tirone M, et al. Early phases of COVID-19 are characterized by a reduction of lymphocyte populations 3 and the presence of atypical monocytes. medRxiv 2020.05.01.20087080; doi: https://doi.org/10.1101/2020.05.01.20087080

12. Laing AG, Lorenc A, Del Barrio ID, Das A, Fish M, Monin L, et al. A consensus Covid-19 immune signature combines immuno-protection with discrete sepsis-like traits associated with poor prognosis. medRxiv 2020 doi: https://doi.org/10.1101/2020.06.08.20125112

13. Organization WH. Clinical management of COVID-19: interim guidance, 27 May 2020. World Health Organization; 2020 Laboratory testing strategy recommendations for COVID-19. Interim guidance. 21 March 2020. Geneva: World Health Organization, 2020. WHO/2019-nCoV/clinical/2020.5

14. Chen X, Ling J, Mo P, Zhang Y, Jiang Q, Ma Z, et al. Restoration of leukomonocyte counts is associated with viral clearance in COVID-19 hospitalized patients. medRxiv 2020.03.03.20030437; doi: https://doi.org/10.1101/2020.03.03.20030437

15. Zheng Y, Huang Z, Ying G, Zhang X, Ye W, Hu Z, et al. Study of the lymphocyte change between COVID-19 and non-COVID-19 pneumonia cases suggesting other factors besides uncontrolled inflammation contributed to multi-organ injury. medRxiv 2020.02.19.20024885; doi: https://doi.org/10.1101/2020.02.19.20024885

16. Tan M, Liu Y, Zhou R, Deng X, Li F, Liang K, et al. Immunopathological characteristics of coronavirus disease 2019 cases in Guangzhou, China. Immunology. 2020; 10.1111/imm.13223. doi:10.1111/imm.13223

17. Wan S, Yi Q, Fan S, Lv J, Zhang X, Guo L, et al. Relationships among lymphocyte subsets, cytokines, and the pulmonary inflammation index in coronavirus (COVID-19) infected patients. Br J Haematol. 2020; 189(3): 428–437. doi:10.1111/bjh.16659

18. Zhou Y, Fu B, Zheng X, Wang D, Zhao C. qi Y, et al. Aberrant pathogenic GM-CSF + T cells and inflammatory CD14 + CD16 + monocytes in severe pulmonary syndrome patients of a new coronavirus. bioRxiv 2020.02.12.945576; doi: https://doi.org/10.1101/2020.02.12.945576

19. Giamarellos-Bourboulis EJ, Netea MG, Rovina N, Akinosoglou K, Antoniadou A, Antonakos N, et al. Complex Immune Dysregulation in COVID-19 Patients with Severe Respiratory Failure. Cell Host Microbe. 2020; 27(6): 992–1000.e3. doi:10.1016/j.chom.2020.04.009

20. Yao XH, Li TY, He ZC, Ping YF, Liu HW, Yu SC, et al. A pathological report of three COVID-19 cases by minimal invasive autopsies. Zhonghua Bing Li Xue Za Zhi. 2020; 49(5): 411–417. doi:10.3760/cma.j.cn112151-20200312-00193

21. Wang K, Chen W, Zhou Y-S, Lian J-Q, Zhang Z, Du P, et al. SARS-CoV-2 invades host cells via a novel route: CD147-spike protein. bioRxiv 2020.03.14.988345; doi: https://doi.org/10.1101/2020.03.14.988345

22. Koch C, Staffler G, Hüttinger R, Hilgert I, Prager E, Černý J, et al. T cell activation-associated epitopes of CD147 in regulation of the T cell response, and their definition by antibody affinity and antigen density. Int Immunol. 1999; 11(5): 777–786. doi:10.1093/intimm/11.5.777

23. Lucas M, Schachterle W, Oberle K, Aichele P, Diefenbach A. Dendritic cells prime natural killer cells by trans-presenting interleukin 15. Immunity. 2007; 26(4): 503–517. doi:10.1016/j.immuni.2007.03.006

24. Cifaldi L, Prencipe G, Caiello I, Bracaglia C, Locatelli F, De Benedetti F, et al. Inhibition of natural killer cell cytotoxicity by interleukin-6: implications for the pathogenesis of macrophage activation syndrome. Arthritis Rheumatol. 2015; 67(11): 3037–3046. doi:10.1002/art.39295

25. Li G, Fan Y, Lai Y, Han T, Li Z, Zhou P, et al. Coronavirus infections and immune responses. J Med Virol. 2020; 92(4): 424–432. doi:10.1002/jmv.25685

26. Cao W, Taylor AK, Biber RE, Davis WG, Kim JH, Reber AJ, et al. Rapid differentiation of monocytes into type I IFN-producing myeloid dendritic cells as an antiviral strategy against influenza virus infection. J Immunol. 2012; 189(5): 2257–2265. doi:10.4049/jimmunol.1200168

27. Spiegel M, Schneider K, Weber F, Weidmann M, Hufert FT. Interaction of severe acute respiratory syndrome-associated coronavirus with dendritic cells. J Gen Virol. 2006; 87(Pt 7): 1953–1960. doi:10.1099/vir.0.81624-0

28. Ziegler T, Matikainen S, Rönkkö E, Österlund P, Sillanpää M, Sirén J, et al. Severe acute respiratory syndrome coronavirus fails to activate cytokine-mediated innate immune responses in cultured human monocyte-derived dendritic cells. J Virol. 2005; 79(21): 13800–13805. doi:10.1128/JVI.79.21.13800-13805.2005

29. Law HK, Cheung CY, Ng HY, Sia SF, Chan YO, Luk W, et al. Chemokine up-regulation in SARS-coronavirus-infected, monocyte-derived human dendritic cells. Blood. 2005; 106(7): 2366–2374. doi:10.1182/blood-2004-10-4166

30. Lau YL, Peiris JS, Law HK. Role of dendritic cells in SARS coronavirus infection. Hong Kong Med J. 2012; 18 Suppl 3: 28–30.

31. Frazier WJ, Hall MW. Immunoparalysis and adverse outcomes from critical illness. Pediatr Clin North Am. 2008; 55(3): 647–xi. doi:10.1016/j.pcl.2008.02.009

32. Wong KL, Tai JJ, Wong WC, Han H, Sem X, Yeap WH, et al. Gene expression profiling reveals the defining features of the classical, intermediate, and nonclassical human monocyte subsets. Blood. 2011; 118(5): e16–e31. doi:10.1182/blood-2010-12-326355

33. Fingerle G, Pforte A, Passlick B, Blumenstein M, Ströbel M, Ziegler-Heitbrock HW. The novel subset of CD14+/CD16+ blood monocytes is expanded in sepsis patients. Blood. 1993; 82(10): 3170–3176.

34. Thieblemont N, Weiss L, Sadeghi HM, Estcourt C, Haeffner-Cavaillon N. CD14lowCD16high: a cytokine-producing monocyte subset which expands during human immunodeficiency virus infection. Eur J Immunol. 1995; 25(12): 3418–3424. doi:10.1002/eji.1830251232

35. Zheng J, Liang H, Xu C, Xu Q, Zhang T, Shen T, et al. An unbalanced PD-L1/CD86 ratio in CD14(++)CD16(+) monocytes is correlated with HCV viremia during chronic HCV infection. Cell Mol Immunol. 2014; 11(3): 294–304. doi:10.1038/cmi.2013.70

36. Azeredo EL, Neves-Souza PC, Alvarenga AR, Reis SR, Torrentes-Carvalho A, Zagne SM, et al. Differential regulation of toll-like receptor-2, toll-like receptor-4, CD16 and human leucocyte antigen-DR on peripheral blood monocytes during mild and severe dengue fever. Immunology. 2010; 130(2): 202–216. doi:10.1111/j.1365-2567.2009.03224.x

37. Hussman JP. Cellular and Molecular Pathways of COVID-19 and Potential Points of Therapeutic Intervention. OSF Preprints. 2020. doi: 10.31219/osf.io/p69g8

38. Taroni JN, Greene CS, Martyanov V, Wood TA, Christmann RB, Farber HW, et al. A novel multi-network approach reveals tissue-specific cellular modulators of fibrosis in systemic sclerosis. Genome Med. 2017; 9(1): 27. Published 2017 Mar 23. doi:10.1186/s13073-017-0417-1

39. Spagnolo P, Balestro E, Aliberti S, Cocconcelli E, Biondini D, Della Casa G, et al. Pulmonary fibrosis secondary to COVID-19: a call to arms? Lancet Respir Med. 2020; S2213–2600(20)30222-8. doi:10.1016/S2213-2600(20)30222-8

